## Supplementary figures and images for "Molecular Matchmakers: Phytoplasma Effector SAP54 Targets MADS-Box Factor SVP to Enhance Attraction of Fecund Female Vectors by Modulating Leaf Responses to Male Presence"

### Figure supplement 1

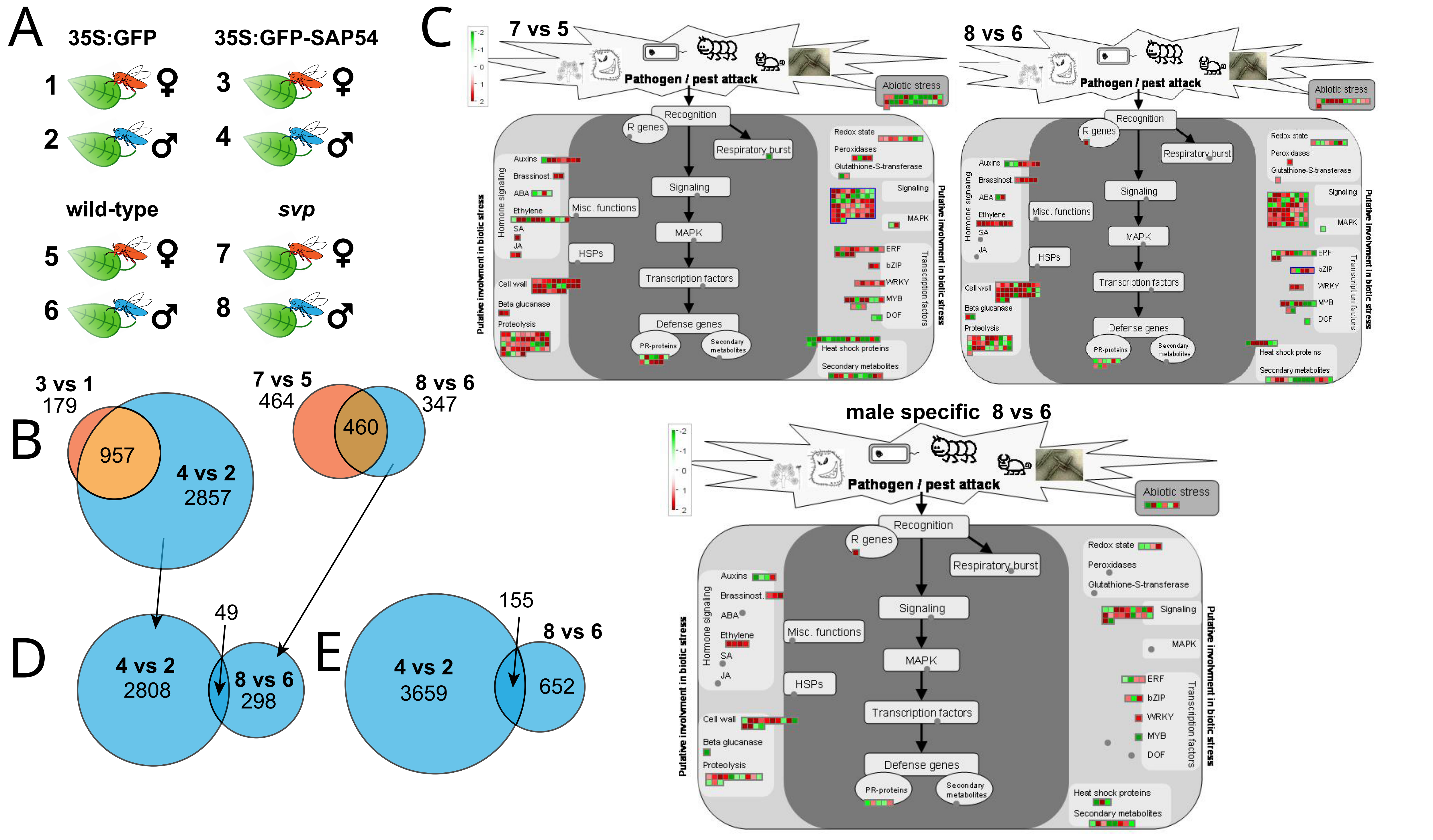

### Figure supplement 2

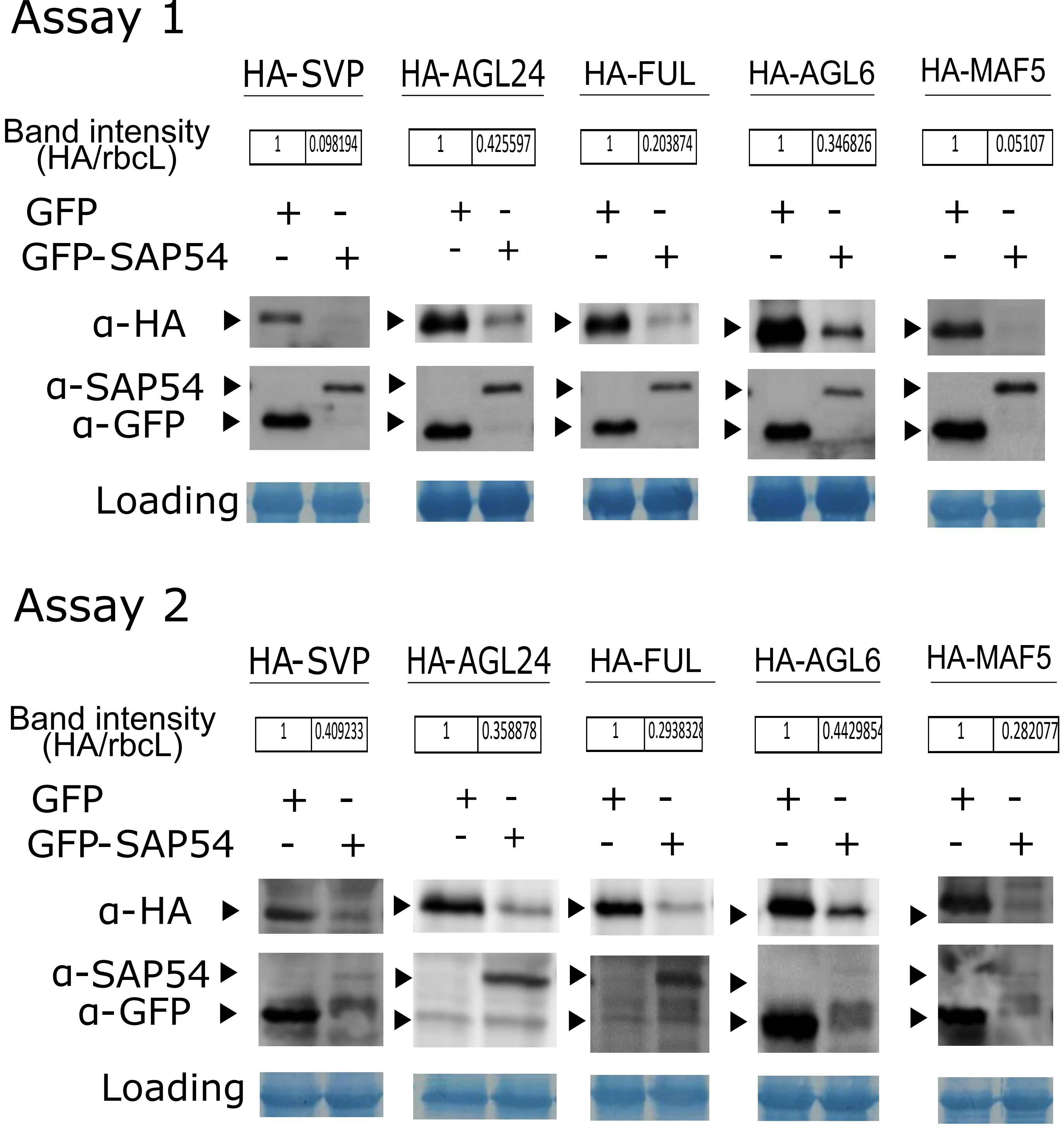

### Figure supplement 4

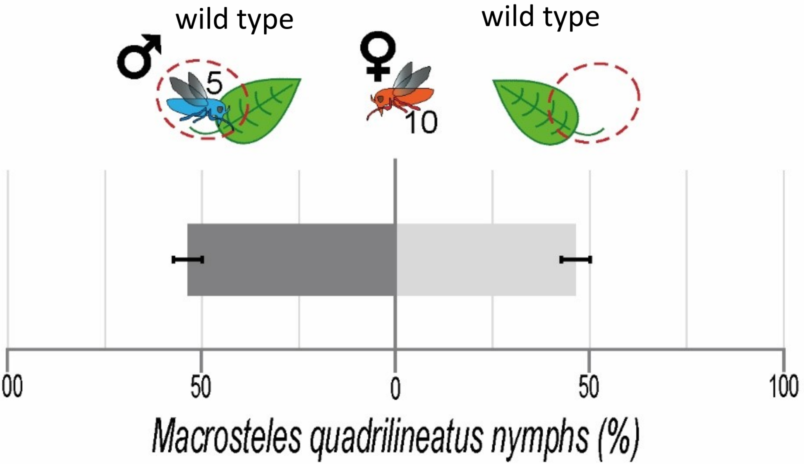

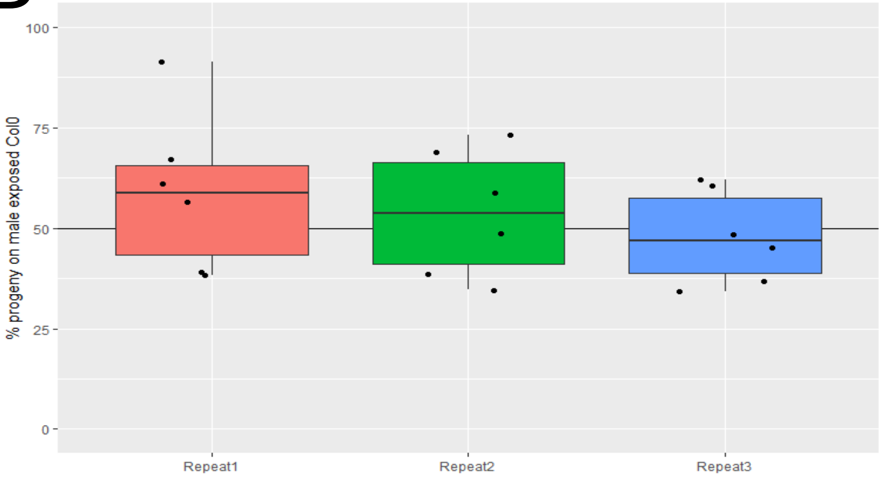

### Figure supplement 5

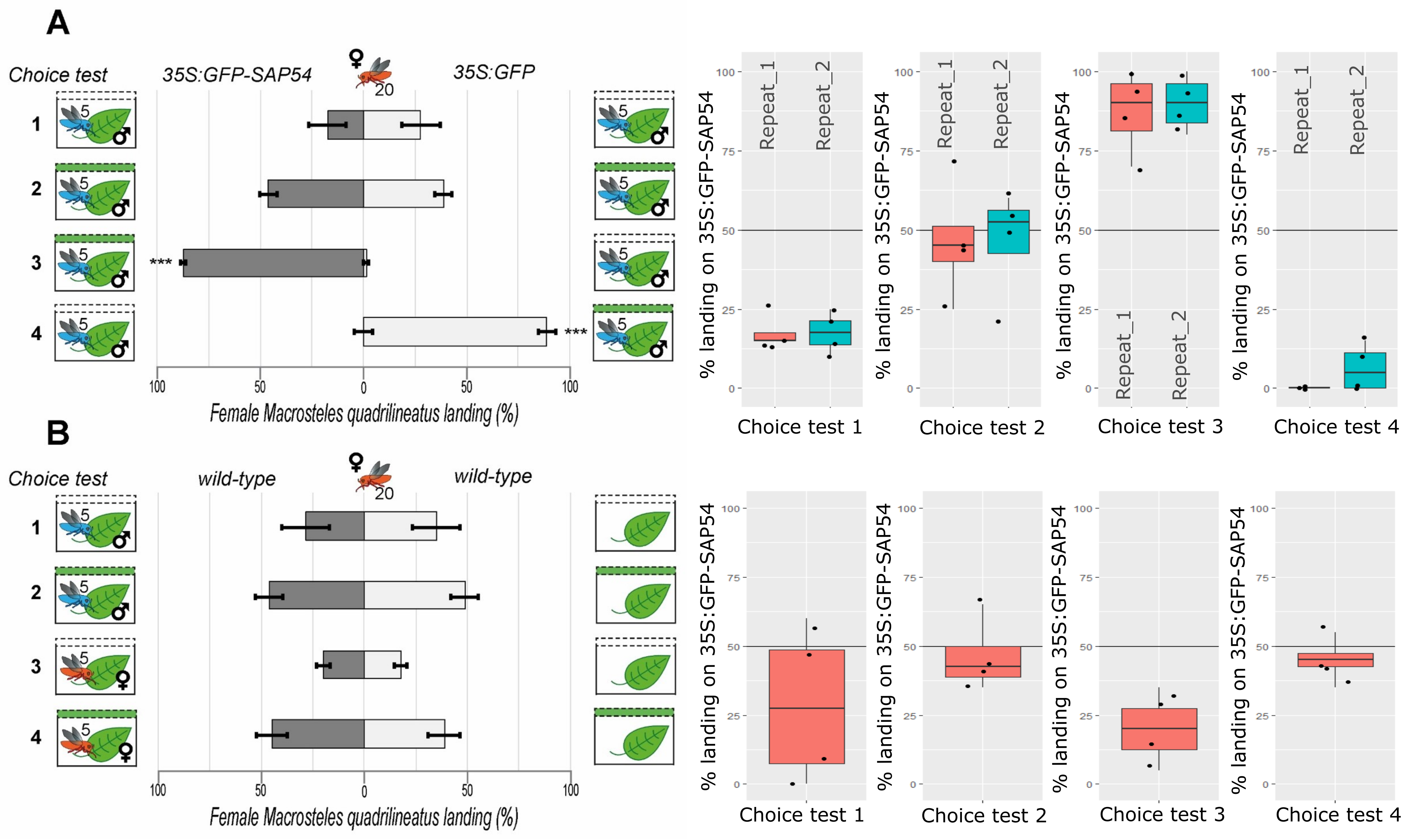

### Figure supplement 6

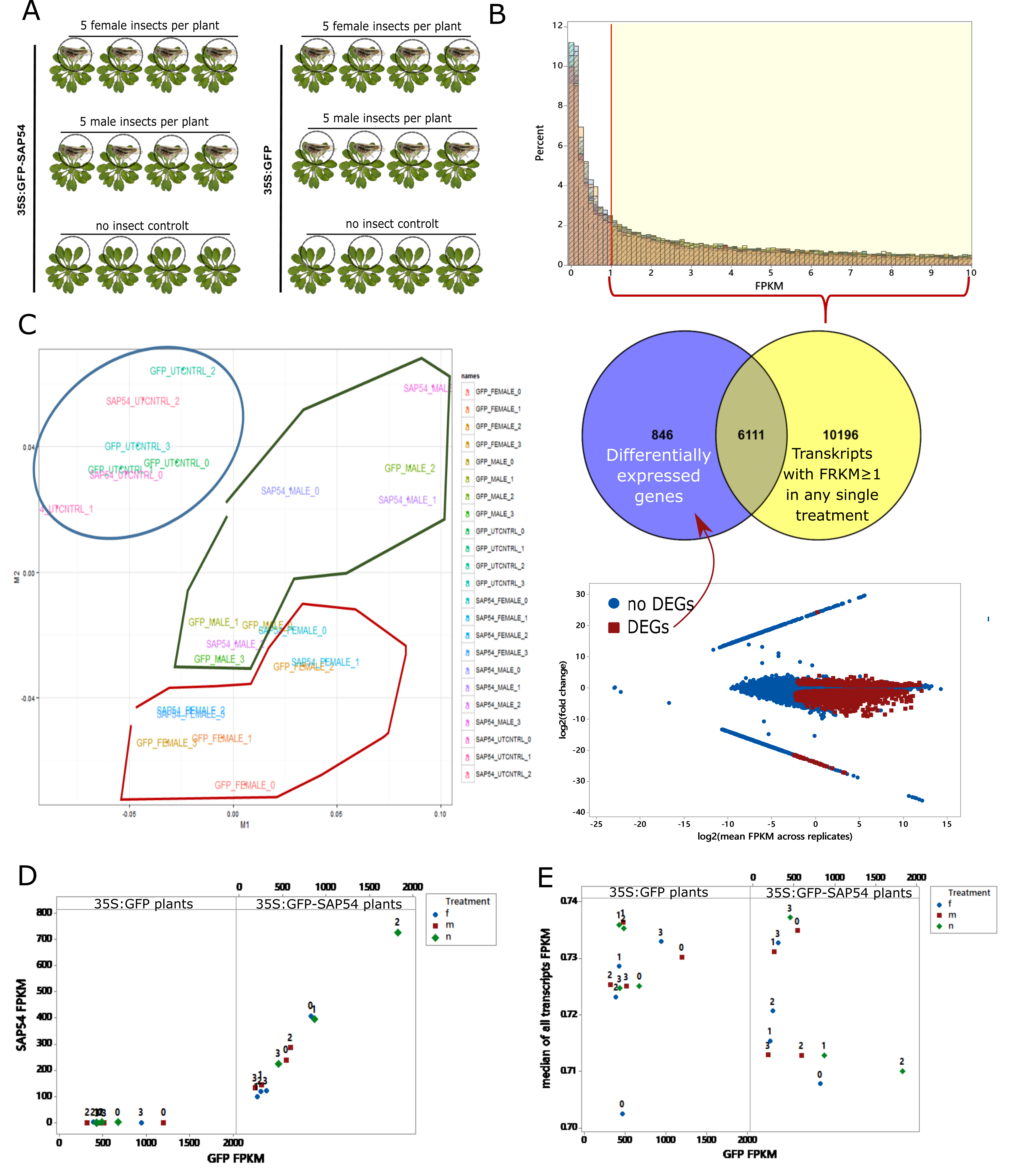

### Figure supplement 7

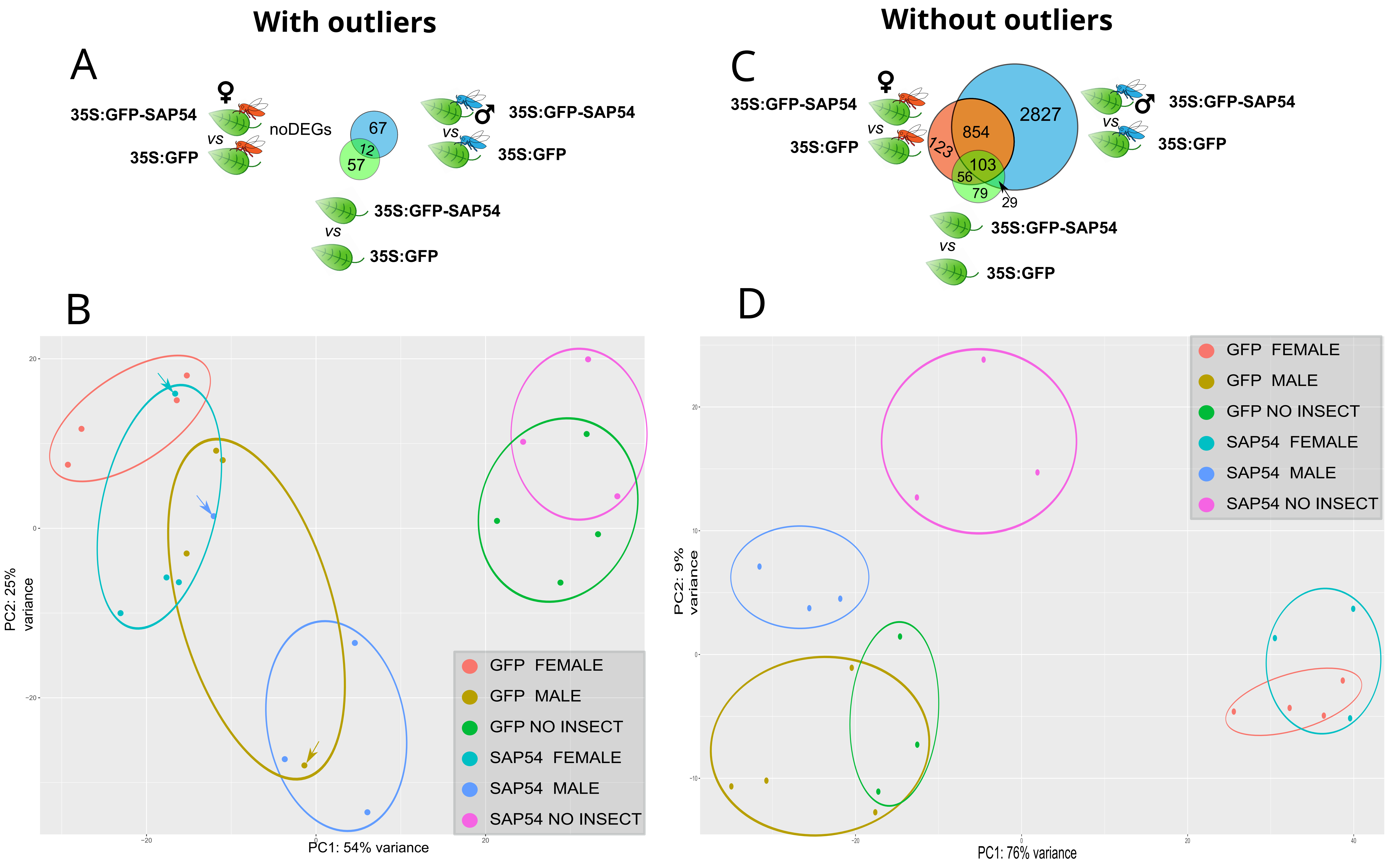

### Figure supplement 8

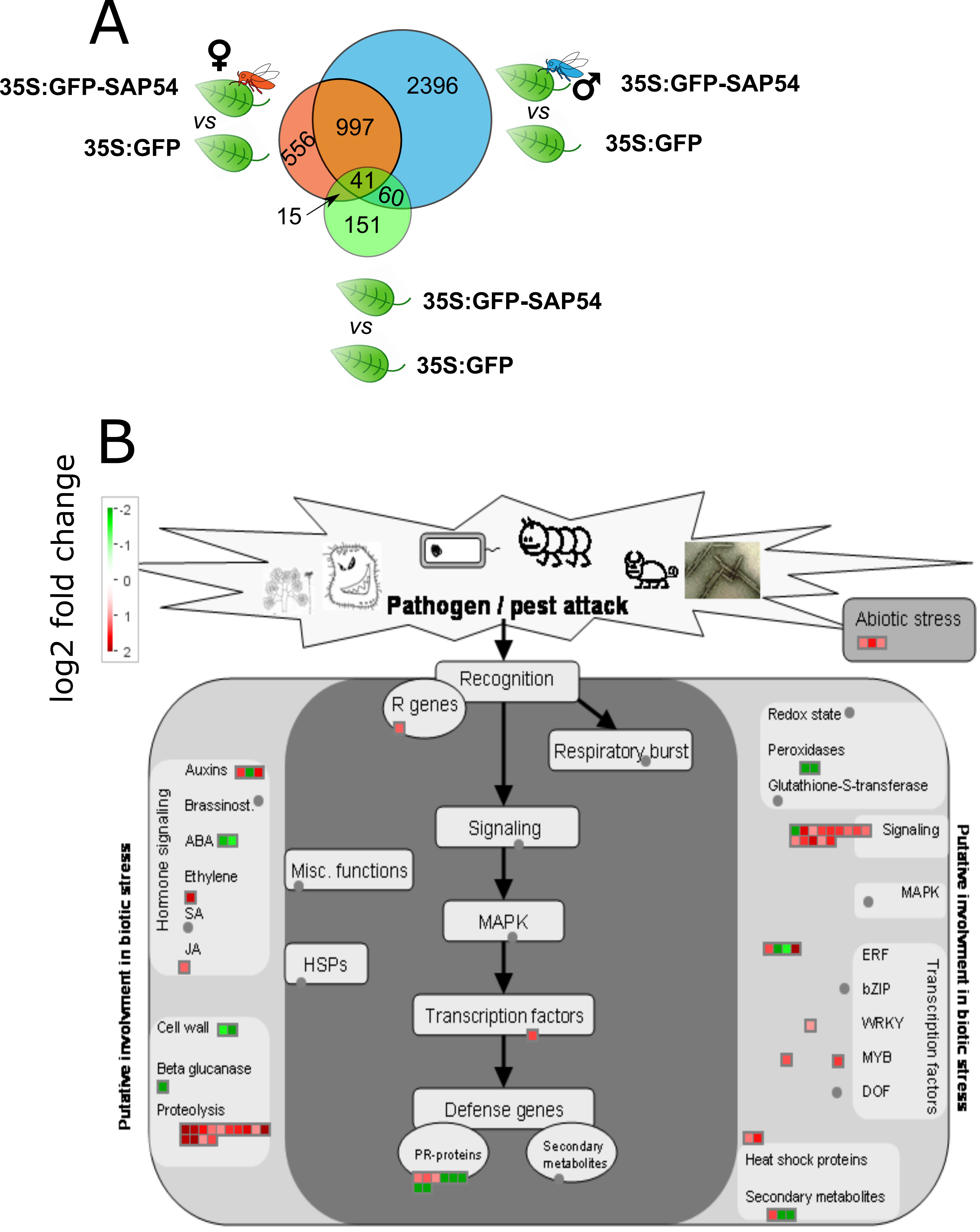

### Figure supplement 9

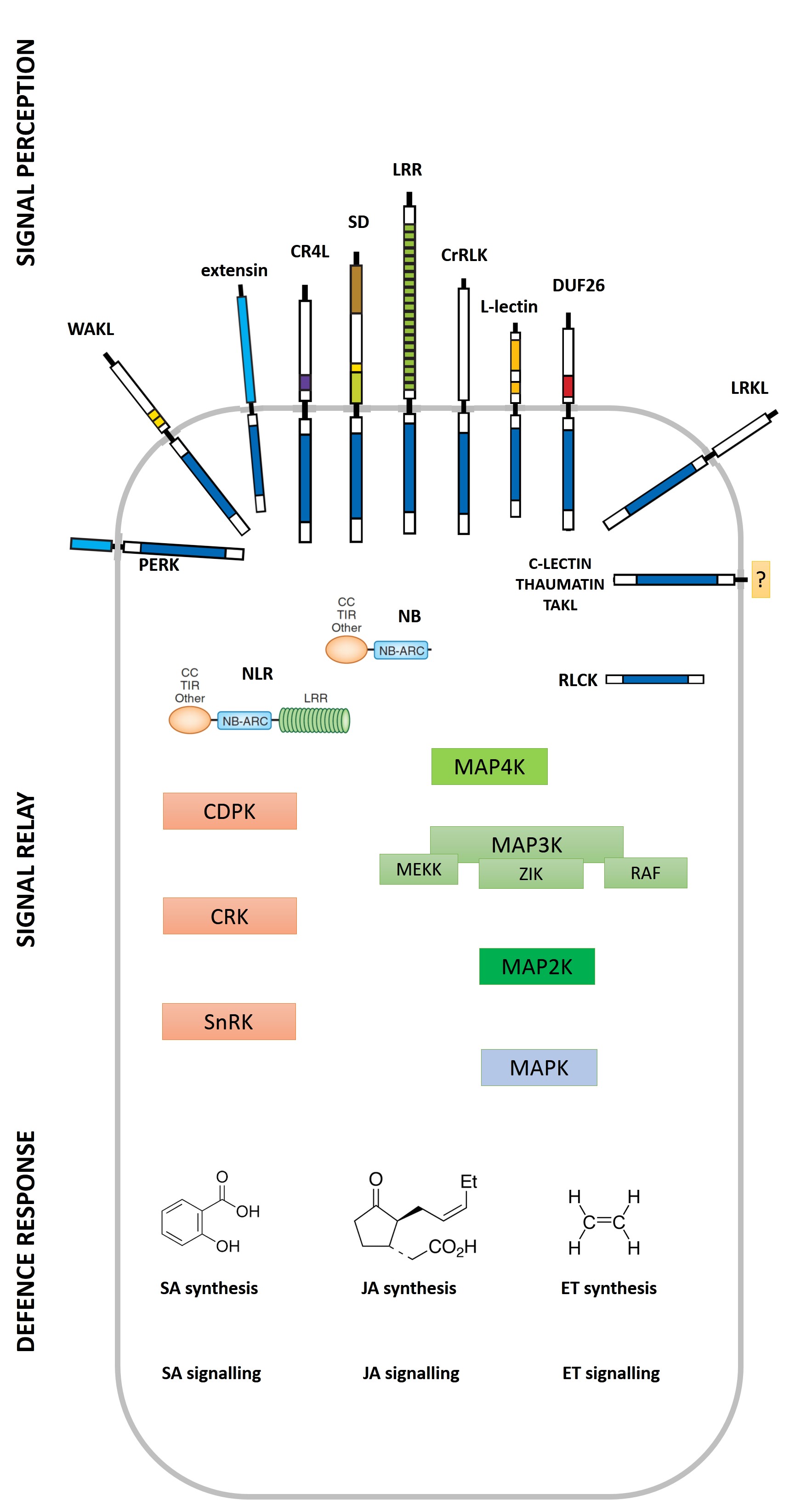
