## Supplementary material for "Molecular Matchmakers: Phytoplasma Effector SAP54 Targets MADS-Box Factor SVP to Enhance Attraction of Fecund Female Vectors by Modulating Leaf Responses to Male Presence": Figure supplement 3

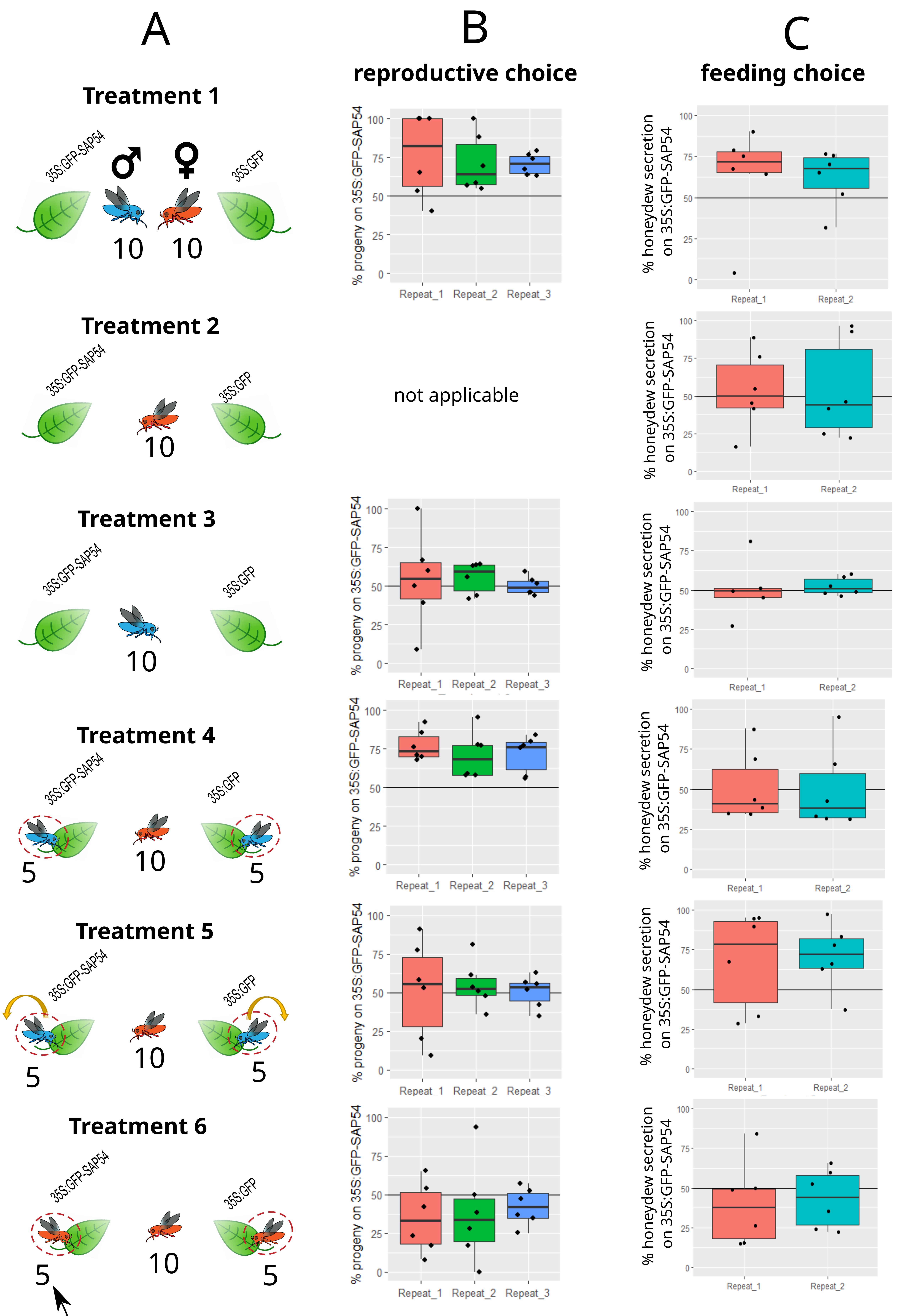


| Treatment | t statistics | p value |
|---|---|---|
| Experiment 1 | 5.2432 | 6.61E-05 |
| Experiment 3 | 0.77136469 | 0.45108 |
| Experiment 4 | 8.04400865 | 3.39E-07 |
| Experiment 5 | 0.49580262 | 0.626382 |
| Experiment 6 | -2.19702566 | 0.042171 |

| Treatment | t statistics | p value |
|---|---|---|
| Experiment 1 | 1.8504 | 0.091275 |
| Experiment 2 | -0.4779467 | 0.642045 |
| Experiment 3 | 0.43710453 | 0.671324 |
| Experiment 4 | 2.69865342 | 0.020707 |
| Experiment 5 | 0.08967685 | 0.930156 |
| Experiment 6 | -0.8572801 | 0.400135 |
