## Supplementary material for "Molecular Matchmakers: Phytoplasma Effector SAP54 Targets MADS-Box Factor SVP to Enhance Attraction of Fecund Female Vectors by Modulating Leaf Responses to Male Presence": Figure supplement 10

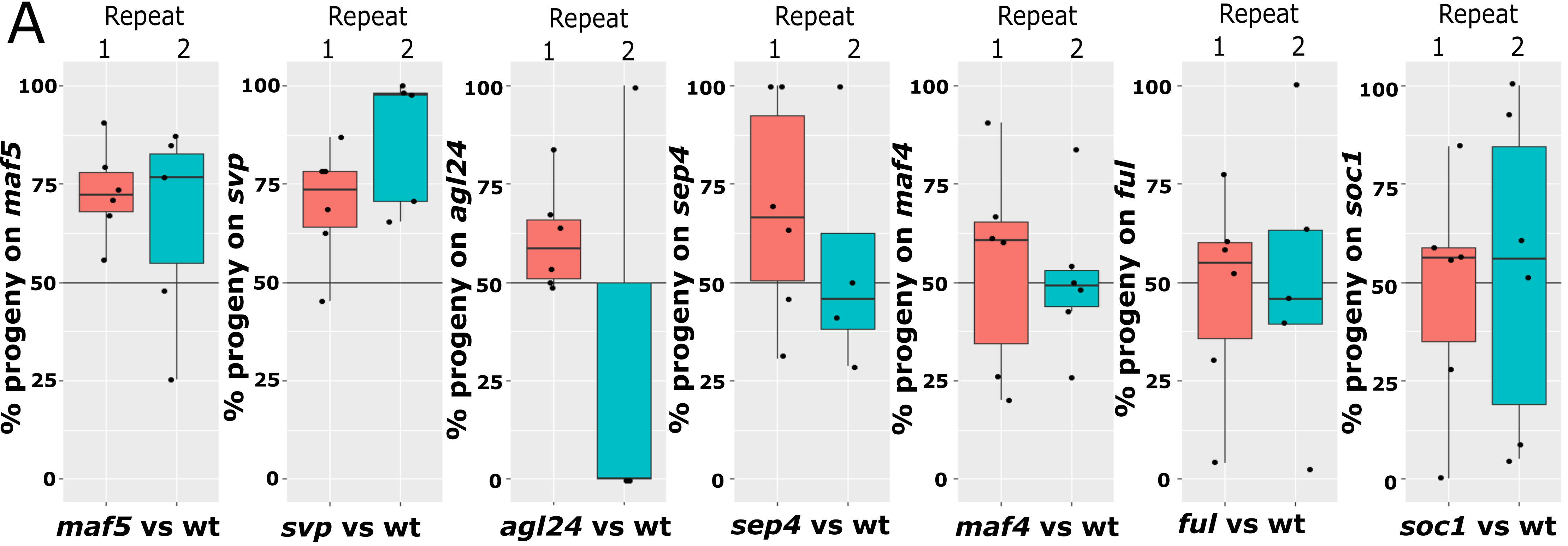

E

| Experiment | test statistics | p value |
|---|---|---|
| maf5 vs wild-type | 3,6430 | **0,0039** |
| svp vs wild-type | 5,2720 | **0,0004** |
| agl24 vs wild-type | 0,1656 | 0,8726 |
| sep4 vs wild-type | 1,4357 | 0,1849 |
| maf4 vs wild-type | 0,3720 | 0,7169 |
| maf1 vs wild-type | 1,2639 | 0,2324 |
| ful vs wild-type | 0,1698 | 0,8686 |
| soc1 vs wild-type | 0,0182 | 0,9858 |
| 35S:SAP54 wt vs 35S:GFP wt | 4,2445 | **0,0004** |
| 35S:SAP54 maf5 vs 35S:GFP maf5 | 4,7577 | **0,0002** |
| 35S:SAP54 svp vs 35S:GFP svp | 1,0168 | 0,3214 |
| maf5 (AYWB) vs wild-type (AYWB) | 0,4514 | 0,6613 |
| svp (AYWB) vs wild-type (AYWB) | 0,0856 | 0,9333 |
| maf5 vs Col0 (female only) | 0,0178 | 0,9861 |
| svp vs Col0 (female only) | 0,6294 | 0,5419 |

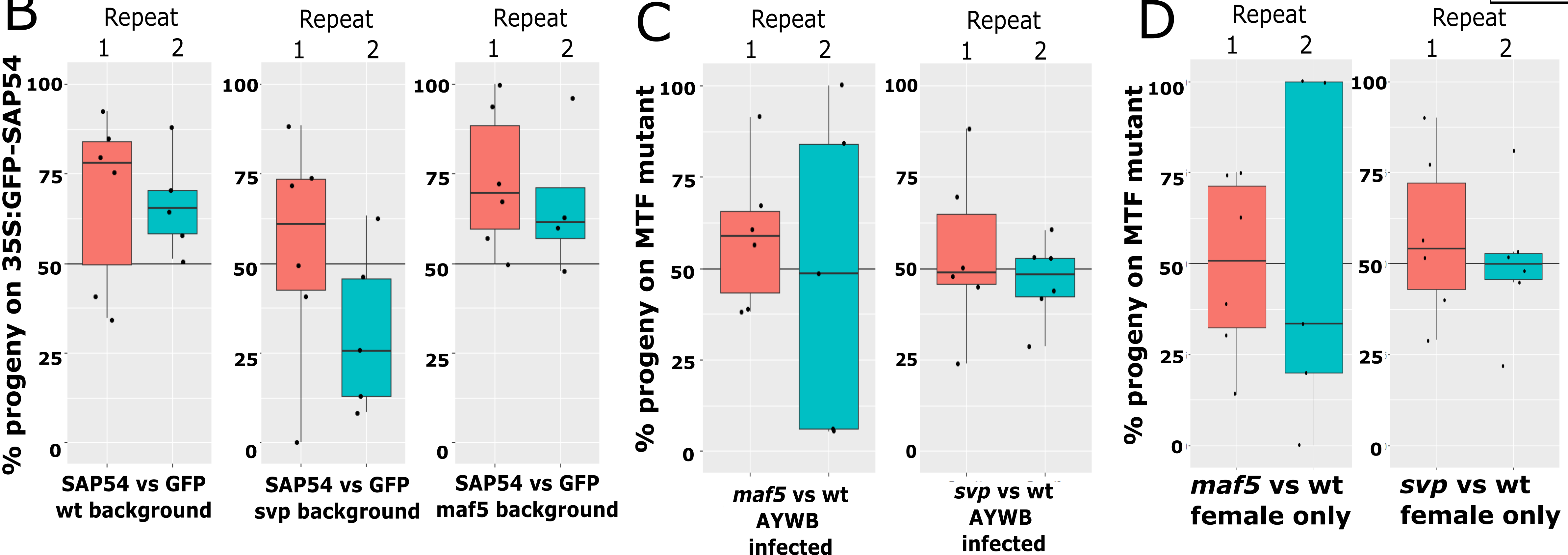
